# Jump: Making Large-Scale Connectomics Accessible

**DOI:** 10.64898/2026.11.12.765999

**Authors:** GP Morris, HF Mohamed, RS Selim, UB Eyo, SK Bonney, A Mughal, S Grubb

## Abstract

Large-scale connectomic resources generated from volumetric electron microscopy (EM) have transformed neuroscience by enabling detailed 3D reconstructions of cellular and subcellular morphology and connectivity across hundreds of thousands of cells in diverse tissues and organisms. Recent advances in machine learning-based cell classification and automated segmentation have enabled large-scale annotation of these datasets, yet barriers remain in validating automated classifications and extracting biological insights. To address these challenges, we developed *Jump*, a suite of user-friendly web-based platforms generated using Claude. These platforms support interactive exploration, expert-guided community review, and quantitative analysis of multiple publicly available EM datasets, while building a growing repository of community-generated annotations. Using µJump, which hosts data from the MICrONS mouse visual cortex dataset, as an example, we demonstrate rapid retrieval of specific cell populations, region-of-interest filtering, biological inspection of cells, and extraction of quantitative morphological and spatial metrics. We showcase analytical features including cell abundance and spatial organisation, connectivity, local cellular neighbourhoods, and volumetric measurements. Using pericytes and endothelial cells as a case study on the cortical vasculature, we further demonstrate how manual review and reclassification can generate accurately identified cell populations for downstream analyses. Consistent with the open-science principles underpinning many large-scale EM resources, Jump is designed to enable broad community participation. By lowering technical barriers to the use of EM datasets, Jump increases analytical efficiency and promotes collaborative efforts to accurately classify cells and structures. Expert-reviewed annotations can be shared across the community, allowing users to build on existing work rather than repeatedly validating the same automated classifications. Collectively, Jump will help researchers to accelerate biological discovery from connectomic datasets.

## Introduction

Recent advances in large-scale electron microscopy (EM), machine learning and computational neuroanatomy have enabled the generation of connectomic datasets at an unprecedented scale. Three leading examples include an entire *Drosophila melanogaster* brain^1, 2^, the MICrONS mouse visual cortex reconstruction^3^ and the h01 human brain dataset^4^. These datasets contain hundreds of thousands of cells and hundreds of millions of synapses, providing unprecedented opportunities to investigate relationships between cellular and subcellular structures, connectivity and function. Such datasets are transforming neuroscience by enabling studies of brain organisation from synaptic morphology to large-scale neuronal network connectivity^5–15^.

To analyse these large-scale EM datasets, researchers have traditionally relied on time-consuming manual efforts^16–21^. However, the rapid growth of connectomic resources has been accompanied by major advances in computational analysis. Automated segmentation and machine-learning approaches now enable large-scale reconstruction and classification of neuronal and non-neuronal cell populations. For example, Elabbady and colleagues developed a hierarchical classifier based on perisomatic ultrastructural features, generating cell-type predictions for tens of thousands of cells throughout the MICrONS volume^13^. In parallel, sophisticated computational platforms have emerged to support the management, annotation and analysis of connectomic data. Connectome Annotation Versioning Engine (CAVE) provides infrastructure for annotation, proofreading, version control and analysis of petascale connectomic datasets, while Neural Decomposition (NEURD) supports automated feature extraction and analysis of neuronal reconstructions^22^.

Despite these advances, using automated classifications to derive biologically meaningful outcomes remains challenging. Many researchers are not formally trained in computational analysis or the complex tools currently required to study EM datasets. Furthermore, biological investigations can be enhanced by enabling researchers to filter specific cells, regions or structures of interest, and importantly, to manually curate errors in automatically classified cells or segmentations, prior to downstream analysis. Furthermore, cells in these volumes are 3D and intertwined yet are usually inspected on flat screens. Tools like immersive virtual reality could therefore enable users to judge depth, appreciate intracellular contacts, trace neuronal morphology^23, 24^ and teach neuroanatomy^25, 26^. However, user-friendly resources to do so on the ultrastructural scale are currently limited. To address these needs, we have developed a suite of tools known as Jump. Our Jump tools are open-source platforms for exploration, validation, segmentation and quantitative analysis of large-scale EM datasets and include a shared virtual reality viewer that runs in a web browser or headset. Here, we describe the design and implementation of one of these tools, µJump, which hosts data from the MICrONS mouse visual cortex dataset^3^. We demonstrate its utility in overcoming barriers to data search and retrieval, data filtering and extraction, and manual cell classification. Although in this introductory work we focus on μJump, our Jump ecosystem currently extends to βJump (a hippocampal volume from a 3xTg Alzheimer’s disease mouse model^9^), δJump (the mouse V1 Deep Dive dataset^15^), ηJump (the human h01 dataset^4^), λJump (Lee16 mouse visual cortex^27^), πJump (MICrONs phase 1 dataset^28^), χJump (a mouse cerebellar cortex dataset^29^), and ωJump (datasets covering 23 tissues from 6 species^30–52^).

## Methods

### Jump Web Application Architecture

µJump and associated tools (δJump, ηJump, βJump, λJump, πJump, χJump and ωJump) are web applications written in HTML, CSS and JavaScript, hosted on GitHub Pages (www.grubblab.com) and version-controlled in a public repository (https://github.com/SorenGrubb/Jump-tools https://doi.org/10.5281/zenodo.24147115). There are many datasets hosted across several tools with links to each available at grubblab.com (Table 1). Each tool uses its source dataset’s own voxel size and units. All code for the web applications was written by a large language model (Claude Sonnet or Opus 4.6-5.5, Anthropic), working from our instructions, an approach often called “vibe coding”. We defined the scientific content, features and design, and did not edit the code by hand. Each change was checked by Claude with automated tests run against a simulated browser, using local copies of the pages and simulated data sources. We also tested each new feature by hand on the live tools to confirm that they worked as intended. The tools were developed iteratively. They began as a simple utility for looking up cells in the MICrONS cubic millimeter dataset, and features were added as new analysis needs arose. Bugs were fixed as they surfaced during use. Development focused on usability, so that the tools could serve both our own and other researchers’ analyses of volume EM data. We also designed them as a learning platform and a community resource.

**Table 1:** Datasets accessible through various tools at grubblab.com.

| Tool | Dataset | Voxel size (nm) |
|---|---|---|
| **βJump** | Alzheimer's vCLEM, hippocampal CA1 | 8 × 8 × 30 |
| **δJump** | V1DD | 9 × 9 × 45 |
| **ηJump** | h01 human cortex | 8 × 8 × 33 |
| **λJump** | Lee16, mouse V1 serial-section TEM | 4 × 4 × 40 |
| **μJump** | MICrONS minnie65 | 4 × 4 × 40 |
| **πJump** | MICrONS pinky100, layer 2/3 | 4 × 4 × 40 |
| **χJump** | Cb2, fragment assembly | 4 × 4 × 40 |
| **ωJump** | 62 volumes — 20 Janelia OpenOrganelle FIB-SEM, 28 served through WEBKNOSSOS, 13 from BossDB and 1 from the Eyewire II consortium; 23 tissues, 6 species | Voxel size column: 4–64 nm (per volume) |

### Data access and visualisation

Image volumes, segmentations and meshes are read directly from the public cloud buckets where each dataset is published, in the Neuroglancer precomputed, graphene and OME-Zarr formats. A coordinate or cell selection is turned into a Neuroglancer state (JSON) and opened in the public viewer with the EM, segmentation and nucleus layers preset. Annotations placed in Neuroglancer, e.g. points (Ctrl+click), lines and polylines, are held in the viewer’s URL, which users may paste back into the tool for it to read the coordinates from, allowing for cells and organelles to be logged without retyping coordinates. Datasets hosted in WEBKNOSSOS are reached through its OME-Zarr streaming interface and shown in Neuroglancer the same way, because WEBKNOSSOS stores annotations on its server rather than in the URL. Cell meshes (multi-resolution, Draco-compressed) are fetched and decoded in the browser, used to compute cell and nucleus volumes, and can be exported as GLB files or as a PowerPoint slide carrying a rotating 3D model.

### Cell nuclei tables to identify individual cells

For each tool a table of nuclei, published cell-type predictions and cortical layer positions was built in advance and embedded in the page, where the dataset allowed it. For minnie65 (µJump; CAVE version 1822) and V1DD (δJump; CAVE version 1196), nucleus detections and cell-type predictions were downloaded from CAVE and joined by nucleus position. For pinky100, the published soma table with its excitatory, inhibitory and glia classes (Zenodo 10.5281/zenodo.3710459) was linked to nucleus IDs by reading the nucleus segmentation at each soma position. For h01, published cell classifications were linked to soma positions via the segment ID at each soma centroid. The Alzheimer’s vCLEM (βJump) and Lee16 (λJump) volumes have no published nucleus table. For vCLEM, nuclei were detected from the Hoechst channel by thresholding and watershed separation in Python, and for Lee16 they were detected directly from the EM images. The cb2 segmentation (χJump) consists of fragments rather than whole cells, so the proofread cells and cell types published by Nguyen et al. (2023)^29^ serve as the starting list, and users assemble further cells from fragments in the tool. ωJump is deliberately built without a nucleus table, because none exists for its FIB-SEM and WEBKNOSSOS volumes. Instead, it has a box sampler that presents randomly selected 20 µm boxes, in which users mark the nuclei they find and name the cells in free text, so the nucleus table is built by the community.

### Cell and organelle tracing directly in the browser

All tools share a module for outlining whole cells, nuclei and organelles directly on the EM. The tracing pad reads image chunks from the dataset’s public volume and shows one section at a time, stepping a user-set interval (five sections by default), so a structure is outlined on a sample of sections rather than on every section. Existing segmentations can be shown over the section for guidance. Outlines are closed either by placing vertices or drawn freehand, freehand strokes being thinned to editable contours. Each structure is typed as a whole cell, a nucleus, or one of 61 organelle and extracellular structure types in a shared ontology, and several structures can be traced on one pad.

Outlining can be assisted by machine learning. Small convolutional networks (U-Nets), exported to ONNX, run in the browser with no server. One clicks inside an organelle to return its outline on that section; a second, trained on the MICrONS nucleus segmentation and hand-traced nuclei, does the same for nuclei. A third network marks candidate organelles on the section, reading the sections 80 nm above and below. Users can accept, reject or relabel each mark, and every answer is stored as a training label for later retraining. An outline can be followed through the stack section by section. Following stops when the area or position changes abruptly, and damaged sections (knife marks, missing tiles) are skipped. When a structure has been drawn by hand on sections 2–25 apart, the sections in between can be filled automatically. Each filled outline blends an interpolation between the two neighbouring hand-drawn outlines with the network’s prediction. The prediction is weighted by how well the network matched the user’s own outlines on those neighbouring sections, so it contributes only where it has proven reliable. Hand-drawn outlines are never altered, and every outline records whether it was drawn, AI-assisted or interpolated.

The volume of each traced structure is updated as the user draws (Cavalieri’s estimator). For each section, the outlined area is multiplied by a thickness equal to half the distance to the section below plus half the distance to the section above. These products are summed over all traced sections. For the first and last traced sections, the missing neighbour is assumed to lie at the same distance as the existing one. The structure therefore extends half a section spacing beyond the outermost outlines, as expected for a structure that tapers off between sampled sections. Contours drawn inside others are subtracted as holes by the even–odd rule. A trapezoid sum over the traced sections only is reported alongside. It cannot include material beyond the outermost contour, so it is a lower bound, and the difference between the two estimates bounds the error at the ends of the stack. A structure outlined on a single section is reported as an area, not a volume.

A surface is lofted in the browser for the 3D preview, optionally rounded between sections by a Catmull–Rom interpolation through four consecutive outlines that affects the preview and the OBJ export but never the volume. Unfinished work is saved as a draft. Upon submission, the outline geometry is written to a JSON file in Drive and the backend sheet gains one index row per tracing - who traced it, what it is, which cell it belongs to, both volume estimates, the section spacing and the file’s URL - with re-shares updating both and the previous version retained. Tracings download per cell as a zip of contours (JSON) and lofted meshes (OBJ, nm), as a single GLB scene in micrometres in which the cell and nucleus are semi-transparent so that the organelles inside are visible, or as a Blender Python script. Both 3D exports place each object at its dataset coordinates, so that several cells opened together keep their true relative positions.

### Google login for community annotation and backend

Users sign in with a Google account (Google Identity Services) and the identity is verified server-side. Identifications, corrections, organelle reports and tracings are stored in Google Sheets through a Google Apps Script web app. Contributors earn points per submission and there are six levels of achievement based on how many points a user accrues, ranging from Novice to Godlike.

### Virtual reality viewer to visualize the 3D structures

Cells selected in any Jump tool can be opened in a shared virtual reality (VR) viewer, a static web page built on three.js and the WebXR Device API. It runs without installation in the browser of a standalone headset, on a PC-tethered headset, or on an ordinary screen. Cell meshes are read with the same in-browser modules as the 3D view and GLB export, so the cells shown are those measured in the tools, in the same micrometre frame. Cells load at a reduced level of detail by default, and only cells on show are loaded at full resolution, to keep the frame rate on standalone headsets. The cells can be placed in context with the cortical layers, a section of the EM, the TriSAM vessel segmentation (Wan & Wei 2025)^53^ coloured by vessel order from our own tracings combined with tracings by the Shih lab^21^, and our tracing of the perivascular spaces. Furthermore, a selected cell’s synaptic partners and touching cells can be brought in.

For excitatory neurons with two-photon recordings (MICrONS functional data, DANDI:000402), the mean calcium response to the repeated movie clips is shown on the cell, matched through the consortium’s manual co-registration, next to the movie the mouse saw. In addition, schematic models built on the real morphologies, synapses and vascular network are provided for teaching: action potential spread, neurovascular coupling^54, 55^, blood flow^56^, sleep^57^, local field potentials^58^, spreading depolarization^59^ and cerebrospinal fluid flow^60, 61^. They are labelled as models in the interface and their parameters and sources are documented in the code.

On a PC, a timeline of camera, view, caption and effect keys turns a scene into a film, which is rendered frame by frame and saved as a video file on the user’s own computer, or as a fisheye dome master for planetarium projection. Scenes can be saved and shared as links under the user’s Google sign-in.

### Manual validation of pericytes and endothelial cells

To demonstrate the utility of μJump for validating and refining automated cell classifications, a region of interest (ROI) was selected that encompassed a 100x100 μm cortical column extending 710 μm down the y-axis of the dataset (start voxel: 143001, 90165, 21647, end voxel: 168001, 267704, 24147). This column was deliberately placed directly adjacent to the MICrONS manually annotated ground-truth region used to train the automated classifier^3, 13, 14^, facilitating manual validation within a highly localised brain region and minimising the potential impact of regional differences in cellular composition on classification performance. It was also designed to replicate the MICrONS ground-truth region dimensions and volume but did not replicate its oblique orientation. All cells detected within the ROI were manually reviewed by a single investigator using μJump. Initial cell-type predictions were obtained from the MICrONS classifications available through CAVE and were assessed against the underlying EM data and 3D reconstructions. Pericytes and endothelial cells were identified based on published characteristics^17, 62, 63^, as described in our previous work^16^. Each cell within the ROI was systematically examined to determine whether it represented a pericyte, endothelial cell, or another cell type, irrespective of its automated classification. For each cell, agreement between the automated and manual classifications was recorded, allowing identification of true-positive, true-negative, false-positive, and false-negative classifications for both pericytes and endothelial cells. Instances where multiple nuclei had been erroneously merged into a single automated detection were identified during manual review and their correct identity was updated prior to analysis. Following manual curation, the corrected classifications were exported from μJump and used to generate confusion matrices and quantify cell-type distributions within the ROI. Nucleus volumes were calculated directly from the decoded nucleus meshes within μJump, as described above, and were subsequently used to compare nuclear volume distributions across cell types in the ROI.

## Results: Tools and Features

### Data search and retrieval

A key feature of µJump design is to enable efficient search and retrieval of cell-specific data from any cell within the MICrONS dataset at the click of a button. This can be achieved in several ways. First, if the coordinate of a specific cell, or subcellular structure of interest is already known in the MICrONS dataset, it can be input into µJump which will then “jump” the user to the midpoint of the cell nearest to that coordinate (Fig. 1A). The identity of the cell will be revealed, alongside key information including which cortical layer the cell resides in, the volumetric centre point of its nucleus, the nucleus and soma volume, the three nearest neighbouring cells and, in some cases, references to published work that have previously studied the cell (Fig. 1B, D-E). Upon clicking the cell name (e.g. “Pericyte” in the example in Fig. 1B), a new tab will open which takes the user directly to the cell in Neuroglancer (Fig. 1C). Alternatively, a user can search cells based on their Root ID, or Nucleus ID, or they can return to previously viewed cells using drop-down menus (Fig. 1A). Finally, we have included an option for users to browse random cells that either have, or have not, been assigned an identity (Fig. 1A). Unclassified cells will be of interest to experienced users who want to identify them, whereas cells that have previously been assigned an identity (either by MICrONS automated cell identification^13^, or by manual user input), are likely to be particularly useful for new users learning how to identify specific cell types in EM datasets or wanting to interrogate specific cell types.

**Figure 1:**
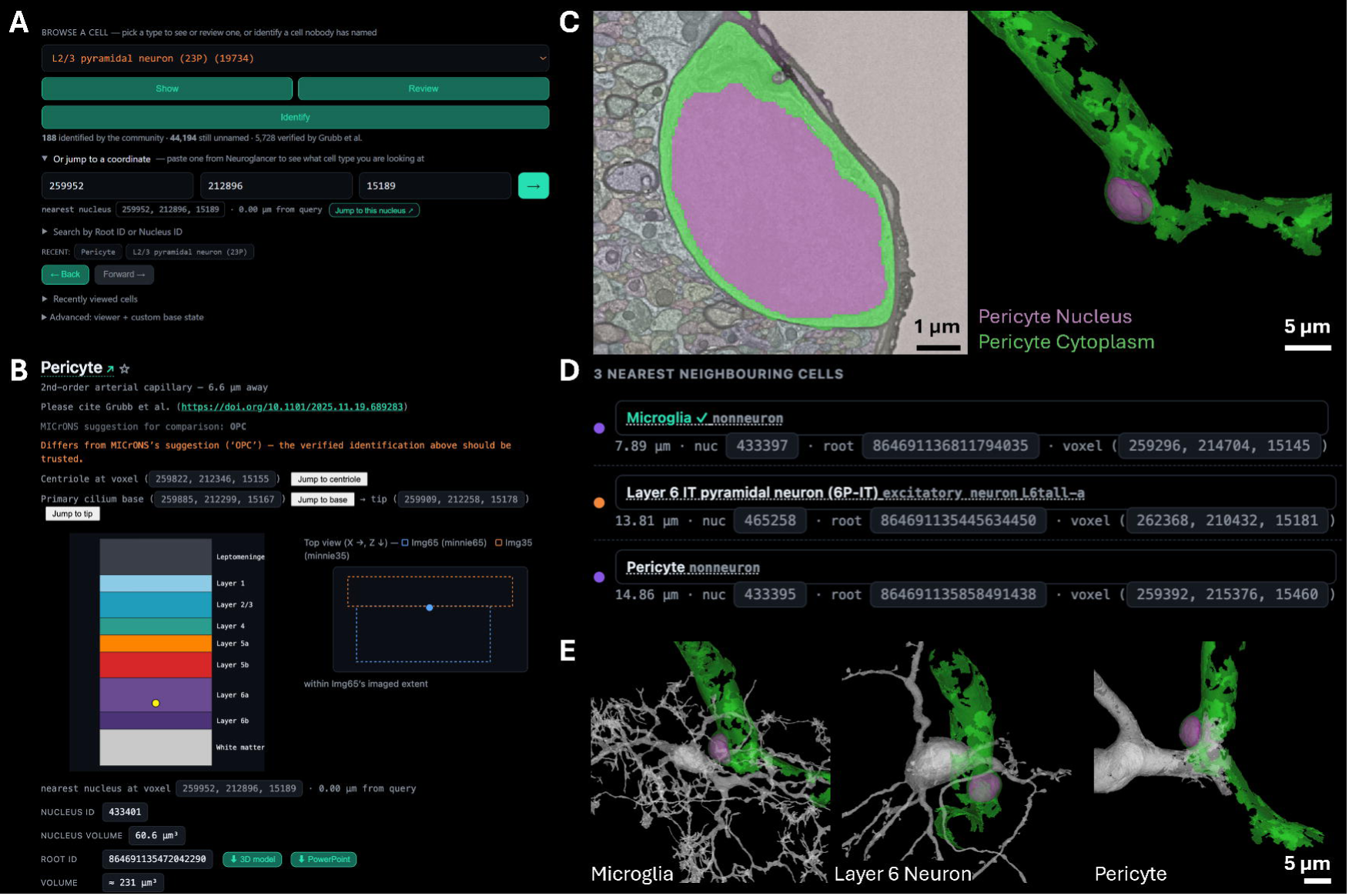
Illustration of µJump interface for cell search, retrieval and navigation. **A:** Search interface showing multiple options for locating cells within the MICrONS dataset, including coordinate-based search, Root ID or Nucleus ID search, navigation to previously viewed cells, and browsing randomly selected identified or unassigned cells. **B:** Cell information panel displaying the identity of the selected cell together with cortical layer location, nucleus midpoint coordinate, nucleus volume, soma volume, and links to external resources for previously identified cells. **C:** Neuroglancer view opened directly from µJump, showing the selected cell in both 2D and 3D with segmented cellular structures (i.e. the nucleus, soma/vessel). The cell segmentations are transparent so that nucleus segmentations are visible. The zoom level and EM contrast are also optimized. **D:** List of the three nearest neighbouring cells based on the relative distance of the volumetric centre point of the cell nucleus to other cell nuclei in its vicinity. **E:** Representative 3D visualisations of the pericyte in B and C and each of its three nearest neighbours.

### Data filtering and extraction

Analysis of large-scale EM datasets has notoriously been a time-consuming process involving manually searching through EM images to find cell types of interest and then manually interrogating those cells to determine various metrics of interest (e.g. nucleus volume, soma volume, contact with other cells and structures, the presence and number of organelles, anatomical position in the dataset and more). We have previously used this approach to interrogate several biological questions in the MICrONS dataset^16–18, 20^. Cell-classification and analyses have, however, been made more efficient by the development of automated tools to segment, classify and analyse 3D reconstructions^3, 4, 12, 13, 22, 64^. Through CAVE, for example, it is possible to download cell-specific information from the MICrONS dataset, including cell-type classifications that were automatically assigned based on perisomatic ultrastructural features^13^. However, there are key issues with this process. First, searching for specific cell types through CAVE to download their information can be cumbersome due to its relatively complex navigation platform, making it challenging for users with limited computational expertise. Second, manually fixing inaccurate automated classifications requires proofreading, but there is no straightforward method for non-experts to reclassify cells or keep a record of re-classifications.

With these hurdles in mind, we designed µJump to provide efficient and straightforward extraction of cell-specific information derived directly from CAVE. For example, one of the key features of μJump is the ability to easily filter the MICrONS dataset by various features, for example cell type, vessel proximity and cortical layer (Fig. 2A). A user can also filter by specific regions of interest by inputting coordinates of a bounding box (Fig. 2B), for example the cortical column used by the MICrONS project used to train their automated classifier^3, 13, 14^, enabling extraction of a variety of data from specific areas. The cell meshes within the bounding boxes can be downloaded as a Google Colab script that makes a Blender animation. The filter can also be applied to organelles annotated in the dataset, so you can query specific information such as “astrocytes carrying lipid droplets in layer 5” (Fig. 2C). Once desired filtering options are selected, data can be visualized in Neuroglancer or downloaded in an Excel file (Fig. 2D). We have also included a live statistics dashboard in a separate tab which contains several summary graphs including the number of community reports over time (with a breakdown on the type of report), the proportion of cells per cell type (broken down by cortical layer), the proportion of classified vs. unclassified cells in MICrONS, nucleus volume (segregated by cell type) and more (Supplementary Figure 1). These features actively update as the manual community reports the identity of cell classifications.

**Figure 2:**
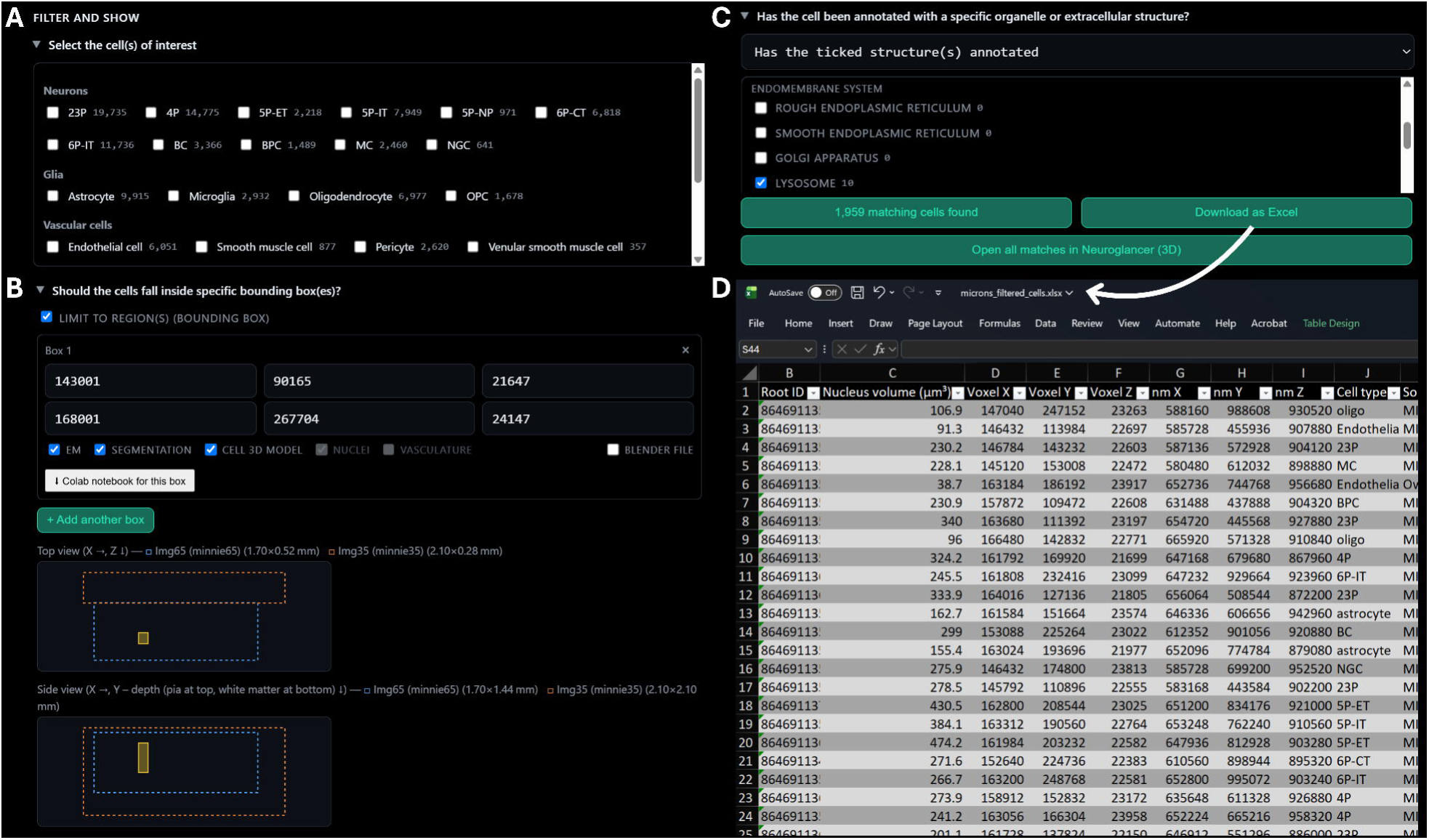
Filtering and data extraction. **A:** Filter interface showing selection of cells within the MICrONS dataset. Note that manual cell type re-classifications within μJump override automatically classified cells derived from CAVE. **B:** Region-of-interest filtering using user-defined bounding box coordinates to restrict analyses to a specified region of tissue. **C:** Organelle and extracellular structure filtering options, enabling selection of manually annotated features including lipid droplets, glycogen, ribosomes, synapses, collagen, extracellular space and pathological structures. **D:** Filtered datasets can be exported directly to a spreadsheet for downstream analysis.

### Manual cell classification and re-classification using guided cell identification

One of the primary drivers to create μJump was to provide a tool to manually reclassify inaccurate automated classifications. To achieve this, we have designed a process called “guided cell identification”. In our Jump tools, manual classifications currently supersede automated classifications, becoming default classifications used for filtering and extracting data. This is based on the premise that manual classifications are currently gold-standard ground truth. However, as automated classifiers become more accurate, this may be revisited in future as they may ultimately be less subjective than a human user.

Guided cell identification has been carefully designed to ensure the classifications can be revised and improved over time, particularly because humans can also make classification errors. In both the case of unclassified cells requiring manual classification (e.g. Fig. 3A), or an automatically classified cell requiring manual re-classification, experienced users who know what the identification of a certain cell type can simply select the correct cell type from a drop-down menu (Fig. 3B). For those learning or new to identifying cell types in EM, they can instead start the process of guided identification (Fig. 3B), which prompts them to answer some simple questions about the ultrastructural features of the cell, “guiding” them towards an accurate classification (Fig. 3C-D). Once the user has determined a cell identity, they are prompted to indicate the certainty of their classification on a five-point scale (Fig. 3E). Once classified, the manually defined identity of the cell (i.e. a “community report”) becomes its default classification within μJump. Importantly, the original automated classification identity is included alongside the manual classification within the μJump interface and remains included in a separate column within downloaded data spreadsheets. Within the online μJump interface, information about which user manually classified the cell is included and users can up- or down-vote these community reports (Fig. 3F) or reclassify a cell if they disagree with a community report. Occasionally, two nuclei are inaccurately segmented together, so there is also an option for users to report dual nuclei and classify the identify of each one separately.

**Figure 3:**
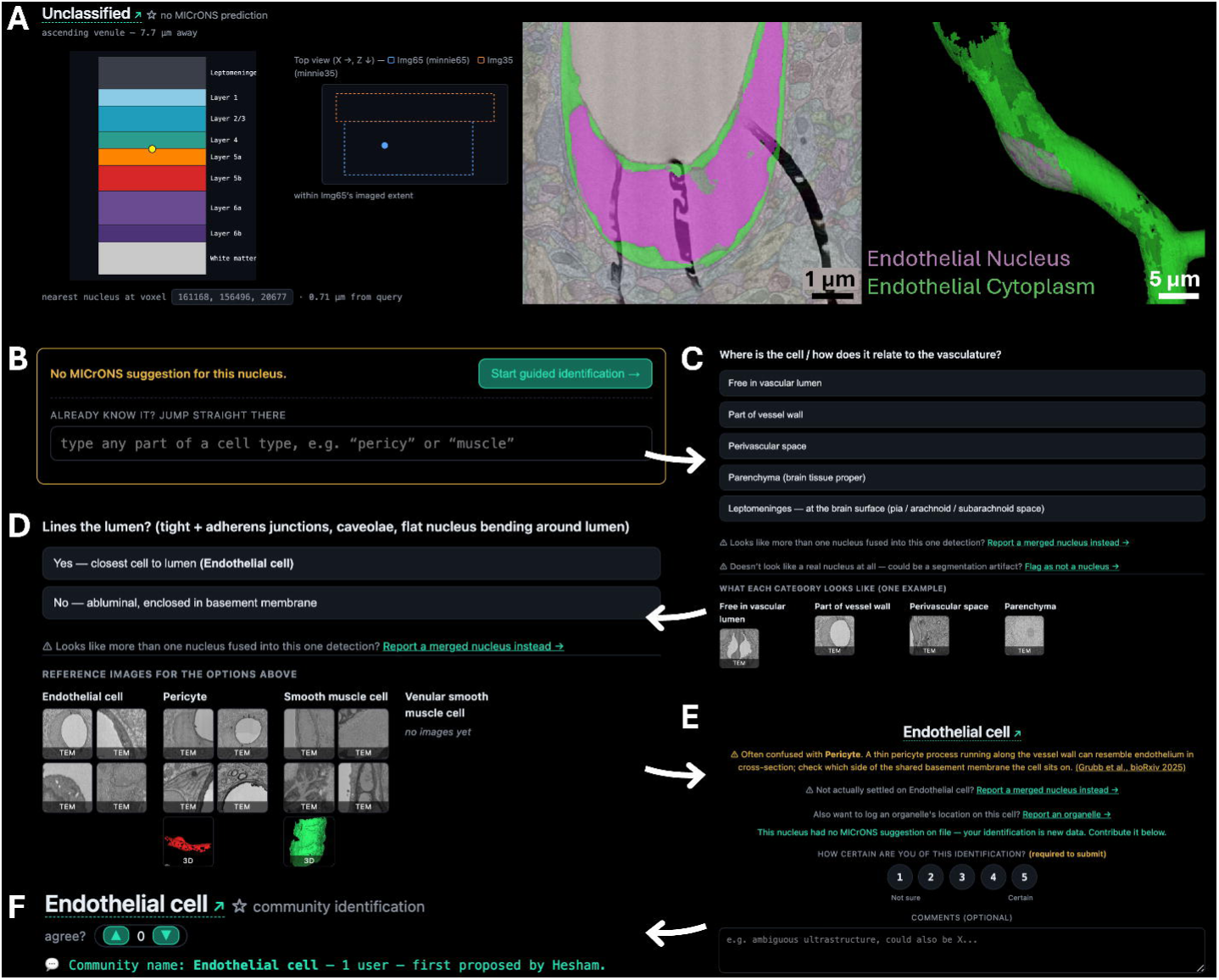
Manual cell classification using guided cell identification in µJump. **A:** Example of an unclassified cell within the MICrONS dataset displayed in Neuroglancer, showing 2D EM imagery and a 3D reconstruction of the segmented structure. A user can use our guided identification process to classify this cell, which in this case is an endothelial cell. **B:** Cell classification interface allowing experienced users to either directly assign a cell identity or inexperienced users to initiate the guided identification process. **C:** First stage of guided identification, in which users answer questions regarding the anatomical location of the cell and its relationship to nearby structures. **D:** Follow-up guided identification step presenting additional questions and representative reference images to assist with distinguishing between candidate cell types. **E:** Final classification screen showing the predicted cell identity based on user responses and allowing users to indicate their confidence in the classification on a five-point scale. **F:** μJump interface displaying manually assigned cell classification from community reports together with information regarding the original classification, who the reporting user was, and options to up-, or down-vote the community report.

To encourage community reports, we have “gamified” the process of guided identification by including a scoring system and leaderboard (Supplementary Figure 2). Users earn points for contributing identifications, confirmations, and other annotations. Accumulated points determine a user’s level (from Novice to Godlike) and position on a leaderboard showing the top contributors. Points and rankings are pooled across the entire Jump ecosystem.

### Connectivity, cell-cell contacts and tripartite synapses

The MICrONS project provides a Connectivity Viewer that enables users to enter the ID of a neuron and explore its synaptic partners. The connectivity data are static and derived from the MICrONS analysis pipeline. They can be viewed either as connectivity graphs or directly in Neuroglancer. We have replicated this functionality in µJump. In addition to displaying the target neuron and its synapses, our implementation also visualizes the segmentations of connected presynaptic and postsynaptic neurons within Neuroglancer (Supplementary Figure 3).

Beyond reproducing the original Connectivity Viewer, µJump introduces a feature that allows users to query contacts between synapses, or a selected cell and other specific cell types. For example, it is possible to identify astrocyte processes located within a user-defined distance of an annotated synapse (Fig. 4A). This provides a rapid way to identify candidate tripartite synapses, where an astrocyte interacts with the pre- and postsynaptic neuronal elements (Fig. 4B-C). Alternatively, cell-cell contacts can be interrogated irrespective of cell type, for example astrocyte-astrocyte contacts (Fig. 4D-F), or any other combination of cell-cell contacts, depending on the user’s interest. The resulting contacts can be explored interactively in Neuroglancer or exported as an Excel spreadsheet for downstream analysis, facilitating targeted investigation of connectivity patterns between specific cell populations, beyond synapses.

**Figure 4:**
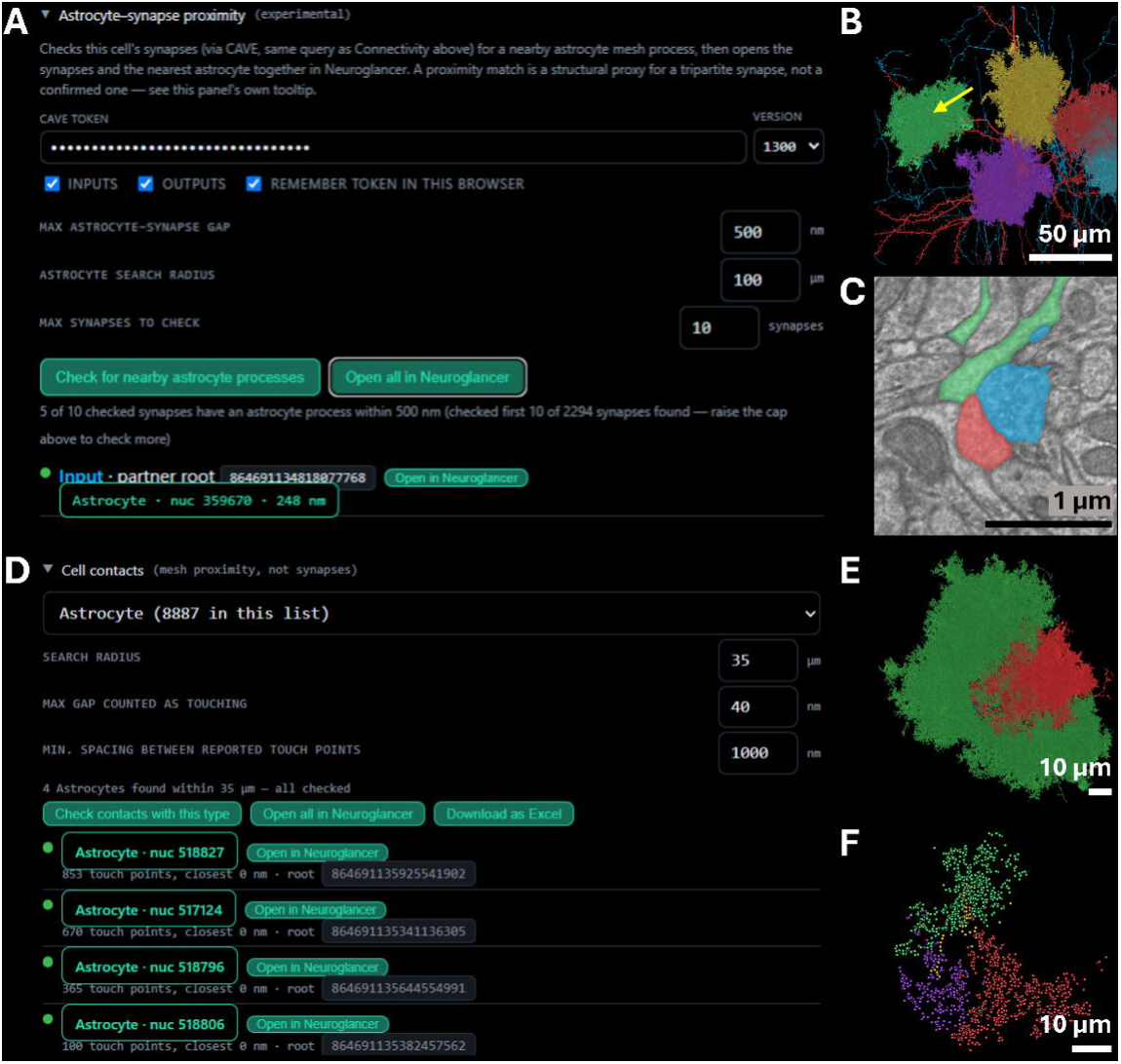
Representative example of astrocyte-synapse and astrocyte-astrocyte proximity data. **A:** Representative example of a search for astrocytes in the proximity of synapses being formed by a single neuron (203360, 133616, 23784), with various options for filtering (astrocyte-synapse gap, astrocyte search radius and max synapses to check). **B:** 3D example of an astrocyte (green) in proximity to a synapse being formed by the queried neuron and an input partner (red and blue, respectively). Yellow arrow indicates the location of EM example in C. **C:** EM example of the synapse between the two neurons (blue and red) and the green astrocyte process in close proximity. **D:** Representative example of a search for astrocyte-astrocyte contacts for a single astrocyte (310304, 116224, 21825), contacting four other astrocytes, with various options for filtering (search radius, max gap counted as touching, min spacing between reported touch points). **E:** 3D example of the queried astrocyte (red) in close proximity to four other astrocytes (all green). **F:** Contact points between the red astrocyte in E and the four other astrocytes (green in E) represented as green, yellow, purple and red annotations.

### Exploring cells in virtual reality

The searches and analyses above all return sets of cells (e.g. a filtered population, the synaptic partners of a neuron, or astrocytes contacting a synapse) whose spatial relationships are often easiest to understand in three dimensions. Any such set can be sent from µJump to a shared virtual reality (VR) viewer, which opens from a single link on a computer screen or in the browser of a VR headset without installation (see Methods). There, the cells can be shown together with the vasculature, cortical layers and the EM itself, and excitatory neurons with two-photon recordings light up with their recorded responses to the movies watched by the mouse imaged by MICrONS^3^. As an example, 26 layer 5 extratelencephalic neurons selected in µJump were opened in the VR viewer and combined with the vasculature, recorded neuronal activity, modelled blood flow, neurovascular coupling, glymphatic flow, simulated sleep, stroke, spreading depression and more into a film using the viewer’s timeline (Supplementary Movie). This scene itself can be explored at https://grubblab.com/vr.html#scene=y3duiire5u.

## Results: Case Study

### A case study of vascular cells to illustrate the utility of μJump

To demonstrate the utility of μJump to assist with improving the accuracy of cell classifications, we manually assessed pericytes and endothelial cells within a defined region-of-interest (ROI) within the MICrONS dataset and compared this to automated classifications derived from CAVE. We deliberately chose to interrogate pericytes and endothelial cells as we were aware MICrONS broad automated classifier did not include a category for endothelial cells^13^, instead often forcing them into the nearest bucket, “Pericyte”, or leaving them “unclassified”, meaning that both pericyte and endothelial cell classifications are often incorrect.

The ROI we analysed contained 1957 automatically classified cells, however we determined that six of these were dual nuclei, meaning there were 1963 cells in the ROI. Within the column we manually identified 77 endothelial cells (3.9% of all cells, Fig. 5A). Due to the lack of an endothelial cell classifier in MICrONS previous work^13^, all of these were falsely negative in the automated classifications, as expected (Fig. 5B-C). These false negatives were largely inaccurately identified as pericytes (51) but some were also unclassified (24) or identified as oligodendrocytes (2). Within the same ROI we found 18 pericytes (0.9% of all cells). Six of these were accurately classified by MICrONS. The remainder were false negative including dual nuclei missed by the automated classifier (6) or classified by MICrONS as either microglia (4) or unclassified (2). There were also 53 false-positive pericytes that were either endothelial cells (51) or smooth muscle cells (2). After manually reclassifying inaccurate classifications, we exported the data from this column and, as an example of the type of data that can be generated, used it to produce an accurate quantification of nucleus volume for every cell type in the ROI (Fig. 5D). In this specific ROI, neurons clearly display the largest nuclei, with mural cells having the smallest (Fig. 5E). We note that these are not the only quantitative data Jump is able to provide. Please refer to Supplementary Table 1 for a full list of quantitative features that can be extracted, noting that these features will be updated over time.

**Figure 5:**
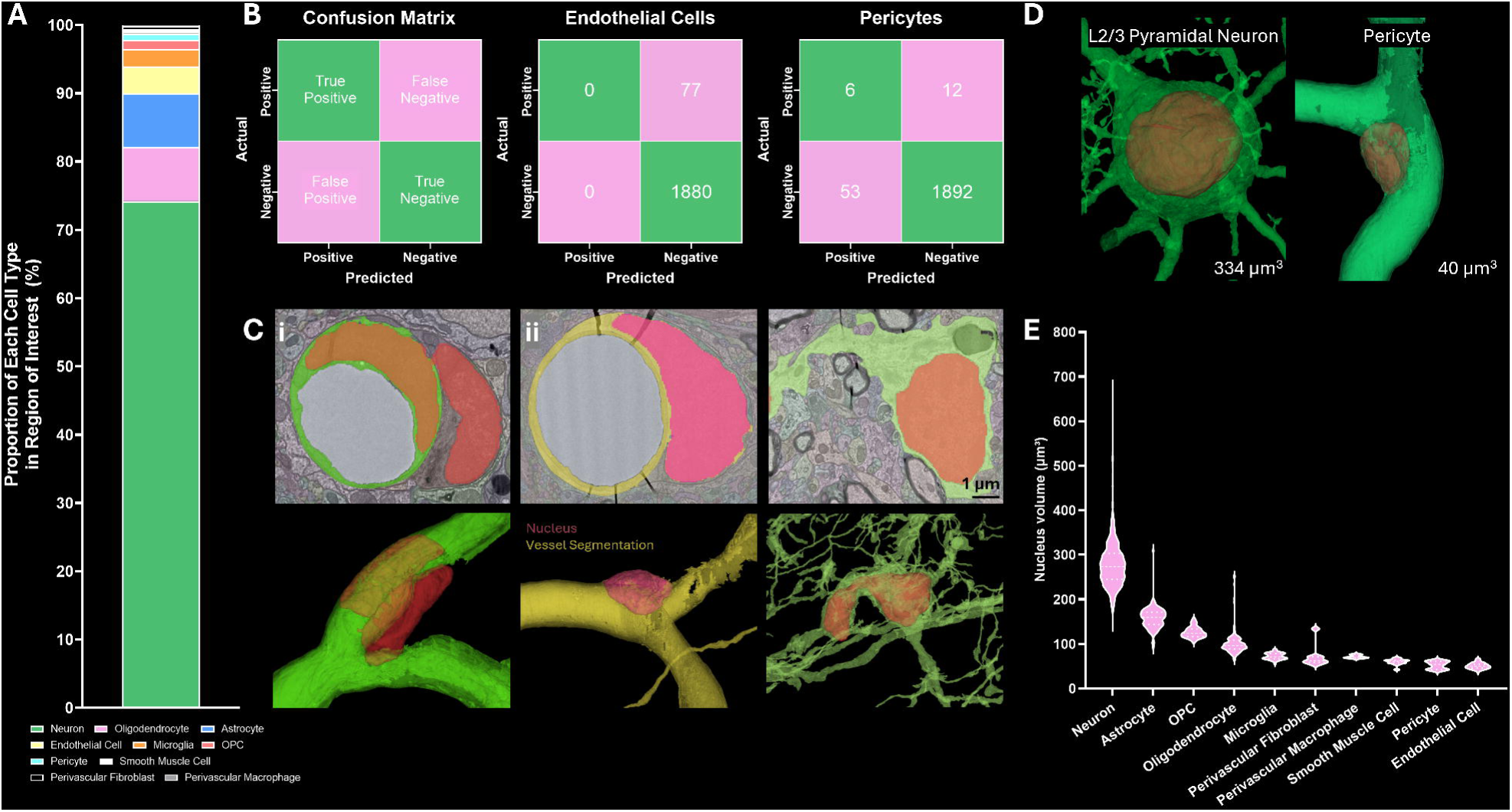
Case study illustrating the capability of µJump to assist with cell classifications and quantification. **A:** Proportion of each cell type in region of interest. **B:** Confusion matrix for endothelial cells and pericytes in region of interest. Note that the endothelial matrix does not add up to 1963 cells because the 6 dual nuclei that were pericytes are not included because they were not classified by MICrONS. **C:** Representative examples of (i) a nuclei (146432, 113984, 22697) classified as a pericyte by MICrONS that is actually a dual nucleus (one a pericyte, one an endothelial cell), (ii) a nucleus (155472, 222176, 21890) classified as a microglia by MICrONS that is actually a pericyte and (iii) a nucleus (157952, 265984, 22178) classified as an OPC by MICrONS that is actually a microglia. Images underneath are 3D reconstructions of the nuclei, vessels and soma from the images above. **D:** Representative examples of the 3D volume of a layer 2/3 pyramidal neuron (left, 164016, 127136, 21805) and a pericyte (right, 161712, 190576, 23900). **E:** Violin plot showing nuclei volume of different cell types. Data is presented with median and upper and lower quartiles (dashed lines within violin plots). The six dual nuclei identified in the ROI were removed from this dataset as their volumes were not truly representative of an individual cell nucleus.

## Discussion

Despite being openly available, large-scale EM datasets remain difficult to utilise for many researchers because effective use of these resources requires expertise in both ultrastructural cell identification and computational analysis. We believe the scientific value of these datasets depends on broad accessibility, as unravelling the vast amount of new biological information they contain will require contributions from many individuals and research groups worldwide. With this goal in mind, we developed the Jump platform to bridge the gap between complex connectomic infrastructure, including key existing tools (e.g. CAVE, Neuroglancer and automated classification pipelines), and the end users seeking to address biological questions. Jump lowers barriers to entry by enabling researchers with limited experience in connectomics or coding to explore, validate and analyse large-scale EM datasets. In this way, Jump complements existing tools by providing a platform for biological interrogation by users with varying levels of computational expertise.

Importantly, Jump reduces the time required for manual interrogation of EM datasets by allowing users to build upon existing automatically generated classifications and community-driven annotations. In reviewing automated classifications we have found many mistakes, for example merged nuclei and cells being inaccurately classified, particularly similar looking cells like OPCs/oligodendrocytes, or pericytes/microglia, which are often mistaken for one another. To overcome this issue, we designed our Jump tools to allow for manual classifications to confirm, or correct automated classifications. Human expertise is particularly important for rare cells, cells with similar ultrastructural characteristics, segmentation errors and classes absent from training data (e.g. endothelial cells, in the case of MICrONS). Manual input is also essential for datasets that have no existing automated classification available. Manual confirmation therefore remains an essential part of the analysis pipeline because classification errors can influence quantitative measures (as demonstrated by our case study of pericytes and endothelial cells), distorting conclusions from downstream analysis. To address the bottleneck of time spent performing manual confirmations, we created a system whereby manual classifications are reported openly, through a community reporting system visible by any user. This allows for individuals to contribute to, and access, community data, negating the need for redundant reclassification of previously identified cells. This is an essential feature of the Jump suite, because no single group could manually assess millions of cells. It is therefore comparable to other citizen science projects like Galaxy Zoo^65^ and Foldit^66^, or, perhaps most similarly, to Eyewire^67^ and Flywire^1^, gamified platforms that were created to crowdsource manual verification of automatically classified neuronal fragments in mouse retina and a whole adult fruit fly brain.

Importantly, in the case of Jump, to maintain reliability, there is a voting system allowing users to up- or down-vote classifications. This does of course open the dataset up to subjective error and bias, so there will be a level of quality control and ongoing revision required over time. For example, users can specifically view community identified cells, allowing experienced researchers to quickly validate other users calls. In essence, this means that publicly available datasets will constantly be evolving as user knowledge evolves, but also as segmentation and annotation tools improve. Ultimately this may mean some biological interpretations could be subject to change, but overall, the accuracy of these interpretations will collectively improve over time.

Beyond improving annotation accuracy, Jump is ultimately designed to facilitate biological discovery. As our Jump community expands, we therefore welcome ideas for improvement, or unique features that can be hosted on the platform. To aid users in proposing analysis tools that could be incorporated into the Jump tools, submitting bug reports, highlight curious features to other users and more, we have provided discussion forums for each Jump platform. One example is our inclusion of trainable classifiers that can be used to speed up the identification and quantification of features (e.g. nuclei, soma, organelles etc) within the datasets. Another, which is unique to our Jump platforms, is the cell-cell contacts feature. Although physical cell-cell contact is not a perfect proxy of functional interactions in the brain or other tissues, in specific contexts it can provide evidence of functional relationships. For example, physical contact is required for cell adhesion (e.g. astrocyte endfeet attaching to cerebrovasculature via receptors through various molecular mechanisms^68^), gap junctions (e.g. for electrical and molecular coupling of neurons and glia^69^), phagocytosis (e.g. microglia clearing pathological proteins, cellular debris and dying cells^70^), and much more. Our cell contact tool is therefore well placed to aid in the discovery of previously underappreciate sites of cell-cell contact that may have important biological implications like, for example, our recent discovery of microglial plugs^16^. A clear limitation is that this tool relies on the accuracy of segmentations, so improvements to segmentation tools will improve the accuracy of this feature over time.

Jump also includes a browser-based VR viewer in which any set of cells or structures collected in the tools can be explored in 3D alongside their vascular and laminar context, and/or with their synaptic partners. Because it runs in a standard web browser, it does not need installation or a dedicated workstation, which will greatly aid use of the tool both for teaching and outreach. Combining the reconstructed cells with functional recordings shows structure and function together, and the same scenes can be turned into films for talks or full-dome projections.

Furthermore, there is clear scope to expand beyond existing datasets, as new datasets become freely available. At present there are eight Jump tools, predominantly hosting brain EM datasets, but also encompassing freely available datasets from 23 different tissues, multiple species and different developmental stages (in ωJump). Importantly, Jump provides a common interface to interact with these various datasets. As the number of publicly available volume EM datasets continues to expand, using Jump as a consistent framework for exploration and analysis will allow users to transfer skills and workflows between datasets regardless of species, tissue type or experimental context.

We note there are a few more important limitations to consider. First, highly active users will likely be contributing to classifications disproportionately, so any systematic errors they make may become widespread. Importantly, data from any user can be removed either partially, or entirely, if such an issue arises. Next, our manual classification process (i.e. guided identification), requires some biological knowledge. To assist users with less expertise in making manual classifications, we have included educational content such as descriptions of certain cells, or example images of various cell types. We have also included the option for every user to indicate how certain they are of a classification, on a 1 to 5 scale. In such cases, an expert review can, in future, validate such classifications. The VR viewer also contains teaching models of activity, blood flow and fluid transport.

In summary, open-access connectomics datasets have greatly expanded in recent years. The key challenge for the scientific community now is to accurately classify cells and structures in these datasets, so that we can correctly interpret them biologically. Jump has been specifically designed to aid in the process, with the overarching goal to make biological discovery within large scale EM datasets easier and more efficient.

## Statements and Declarations

### Author Contribution information

**Gary P. Morris:** conceptualization; data curation; formal analysis; investigation; methodology; project administration; resources; software; supervision; validation; visualisation; writing – original draft; writing – review and editing. **Hesham F. Mohamed:** conceptualization; data curation; methodology; validation; visualisation; writing – original draft; writing – review and editing. **Rukaya S. Selim:** conceptualization; data curation; methodology; validation; visualisation; writing – original draft; writing – review and editing. **Ukpong B. Eyo:** conceptualization; resources; supervision; writing – review and editing. **Stephanie K. Bonney:** data curation; investigation; software; validation; writing – review and editing. **Amreen Mughal:** data curation; investigation; software; validation; writing – review and editing. **Søren Grubb:** conceptualization; data curation; formal analysis; investigation; methodology; project administration; resources; software; supervision; validation; visualisation; writing – original draft; writing – review and editing.

## Acknowledgments

We want to thank all the individuals behind the datasets we have used in this study. If these datasets had not been made public this work would not have been possible. We also thank Vanshika Chaddha for manual annotation of vascular cells during her summer internship at the National Institute of Health. Microsoft Copilot and Claude were used during manuscript preparation to assist with language refinement, grammatical correction, and improving clarity of expression. These tools were not used to generate scientific interpretations or develop the intellectual content of the manuscript. All content assisted by these tools was reviewed, verified, and edited by the authors, who take full responsibility for the accuracy, interpretation, and integrity of the manuscript.

## Data Availability

All software is available at the links found within the paper. All data supporting the findings of this study are available within the paper, its supplementary information, or our Jump platforms.

