## Supplementary Materials for "Jump: Making Large-Scale Connectomics Accessible"

Dr. Gary Morris

Full address: Medical Sciences Precinct, 17 Liverpool Street, Hobart, TAS 7000, Australia, Ph: +61 3 6226 4851;

**Or:**

Dr. Søren Grubb

Full address: Blegdamsvej 3B, Copenhagen N, Denmark, DK-2200, Ph: +45 27290007,

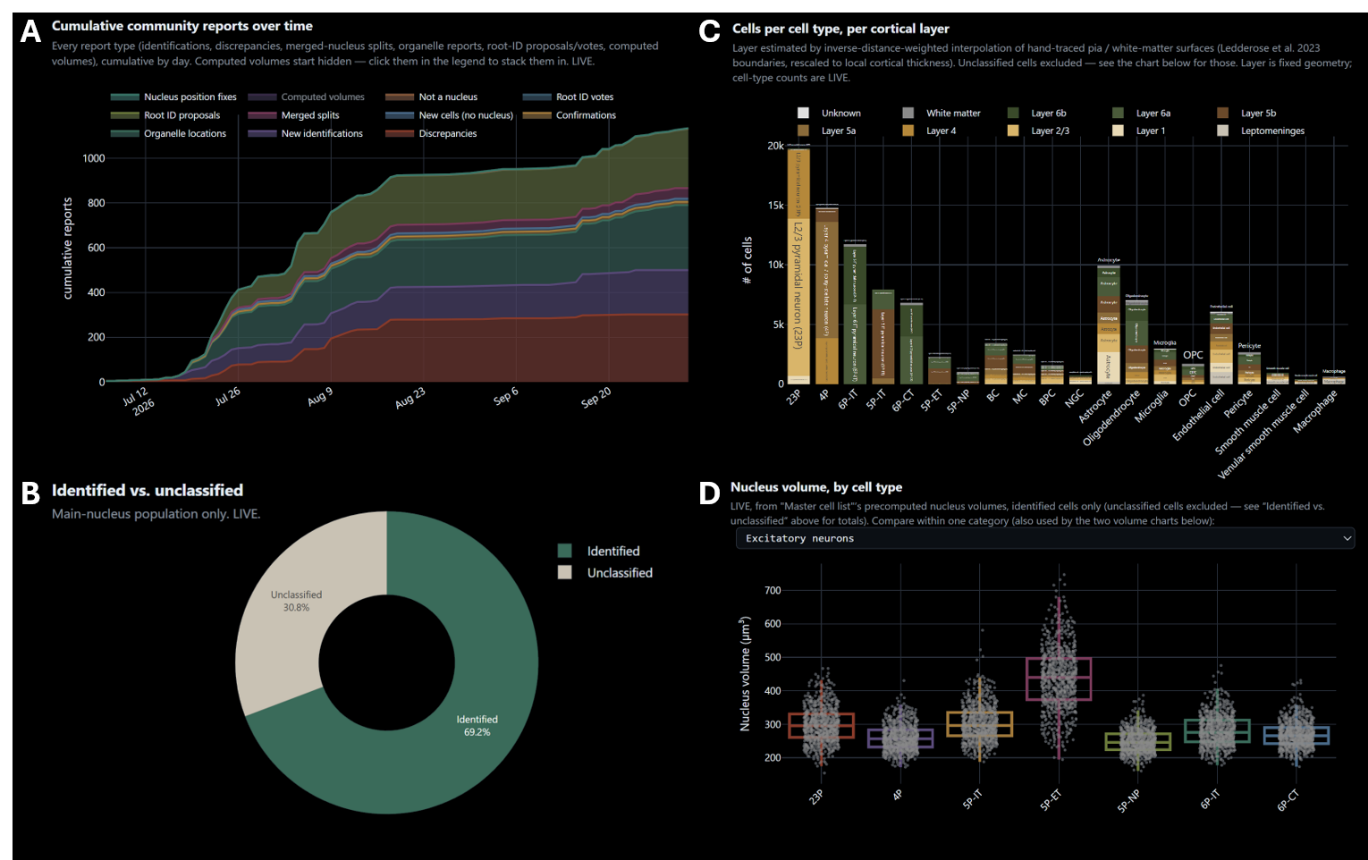

**Supplementary Figure 1: Examples of data included in the live statistics dashboard. A:** Community reporting dashboard showing cumulative manual classifications and annotations submitted over time. **B:** Proportion of cells either classified (e.g. identified) or unclassified in µJump. **C:** Distribution of classified cells by cell type and cortical layer. **D:** Nucleus volumes segregated by cell type.

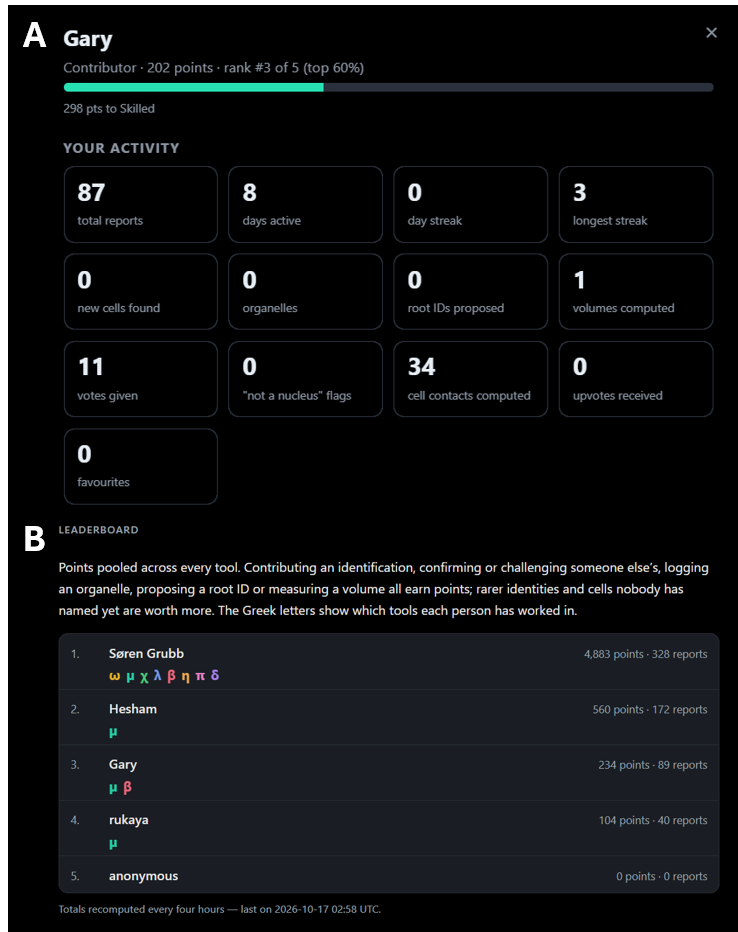

**Supplementary Figure 2: Example of scoring system and leaderboard. A:** Representative example of activity by a single user in µJump. **B:** Leaderboard hosted at grubblab.com showing pooled points from across every Jump tool.

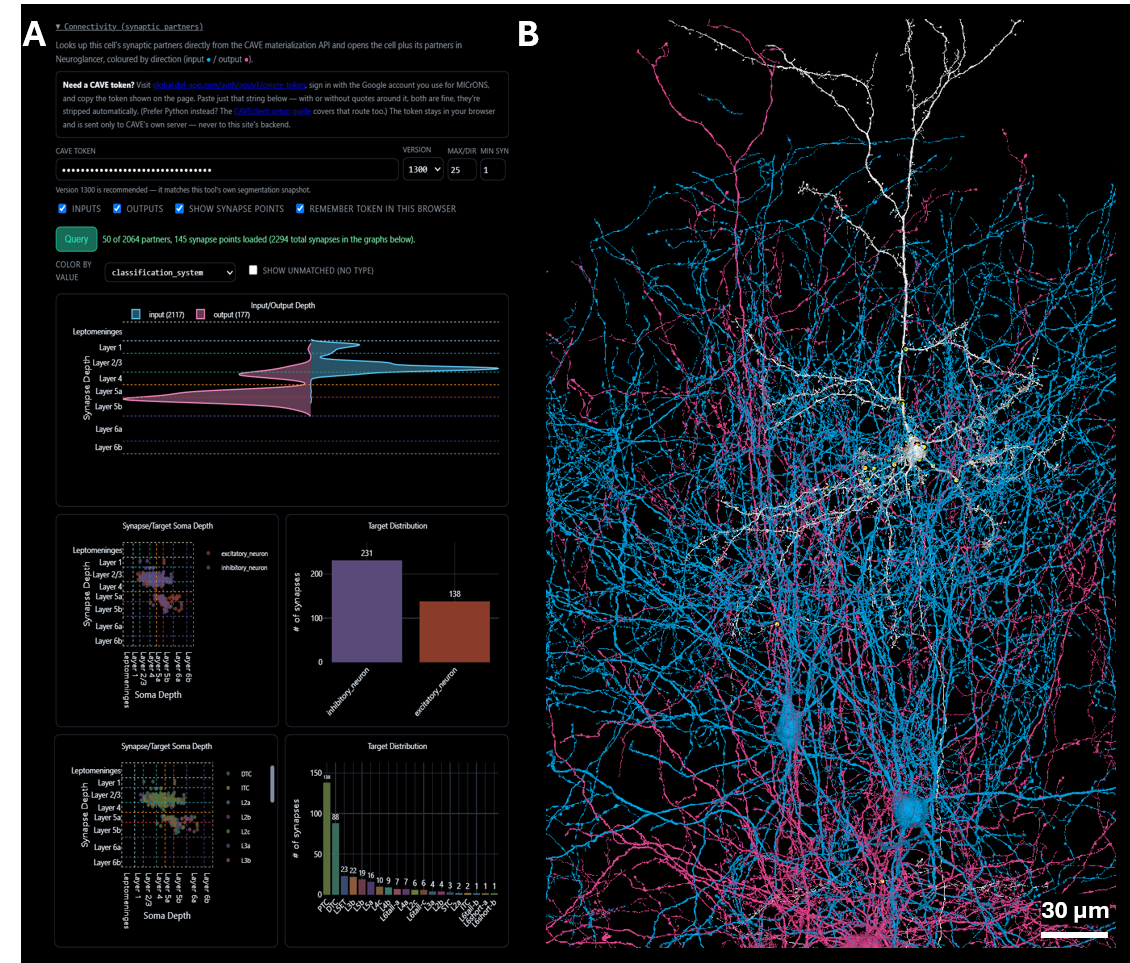

**Supplementary Figure 3: Representative example of a neuron, its synapses and segmentations of synaptic partners. A:** Representative example of connectivity data for a single neuron (203360, 133616, 23784) with µJump. **B:** 3D reconstruction of the neuron in A (white), synaptic contacts (yellow annotations) and synaptic partners (blue and red).

### **Supplementary Table S1. Features of µJump**

Features are grouped by the stage of analysis they serve: finding a cell (A), reading its context (B), measuring it (C), identifying and curating it (D–F), analysing populations (G), connectivity and overview (H), data provenance (I), and machine-learning assistance (J).

| **#** | **Feature** | **What it does** | **How it helps analysis** |
| --- | --- | --- | --- |
| A1 | **Coordinate jump** | Takes x, y, z in voxels and opens Neuroglancer centred on that point, with EM, cell segmentation and nucleus layers | Any coordinate from a paper, table or colleague can be revisited in one step, and the shared link reproduces the same view. |
| A2 | **Nearest-nucleus lookup** | Resolves the coordinate to the nearest of 144,118 automatically detected nuclei and reports nucleus ID, cell root ID, predicted cell type and distance. Both IDs are pre-selected in the viewer. | Links a point of interest (a synapse, a vessel feature, an organelle) to the cell it belongs to, without a CAVE account. |
| A3 | **Neighbouring cells** | Lists the three nearest cells with type and distance. One click moves to a neighbour; “Back to previous cell” returns. | Quick survey of a cell’s immediate surroundings, e.g. satellite glia or perivascular cells. |
| A4 | **Random example of a cell type** | Loads a randomly chosen cell of the selected type. Generic fallback labels are excluded, and the chosen type is kept between draws. | Unbiased spot-checks of a class, and practice material for learning the identification criteria. |
| A5 | **Previous / Next of the same type** | A row on the identity card walks every cell with the same identity in a fixed order, with a counter (e.g. “Astrocyte · 3 of 57”). | Complete, systematic review of one cell class. |
| A6 | **Tabs and deep links** | Jump, Filter and show, References, Discussion and Statistics tabs. | Inspection, population analysis and discussion in one window. |
| B1 | **Layered identity** | Shows the cell’s identity and its source, in order of precedence: expert-verified, community consensus, then the MICrONS model prediction (broad cell class and fine morphological type). | The provenance of each label is visible, so predictions can be used where reliable and challenged where not. |
| B2 | **Expert-verified reference set** | About 14,300 expert identifications, matched to detected nuclei within 5 µm, override the model prediction. Includes cells without a detected nucleus and nuclei merged by the detector. | Ground truth for estimating prediction accuracy and for scoring annotators. |
| B3 | **Cortical depth and layer** | Estimates layer from the cell’s position between pia and white matter (inverse-distance weighting over 24,790 pial and 2,630 white-matter traced points). Layer boundaries are rescaled for aldehyde-fixation shrinkage. Drawn in a layer model and a top view. | Laminar position for every cell, including cells with no model prediction; enables layer-resolved comparisons. |
| B4 | **V1 column membership** | Flags cells inside the MICrONS V1 column (1,899 cells); the column boundary can be drawn in the viewer. | Links any cell to the most densely proofread and classified subvolume. |
| B5 | **Visual area** | Assigns each cell to VISp, VISrl, VISal or VISlm from the published area-boundary meshes, evaluated at the cell’s own depth. | Area-specific comparisons of cell types and organelles. |
| B6 | **Distance to vasculature** | Distance from each nucleus to the nearest vessel (k-d tree, no distance cap) and that vessel’s class among 10 vessel type/order categories. | Separates vessel-associated cells (pericytes, smooth muscle cells, perivascular macrophages and fibroblasts) from parenchymal cells; supports neurovascular analyses. |
| B7 | **Distance to glia limitans and white-matter border** | Distance from each nucleus to the pial glia limitans and to the white-matter border. | Identifies surface- and border-associated populations, e.g. glia limitans astrocytes. |
| B8 | **EM section beside the model** | One EM plane through the nucleus (32 nm per pixel, about 8 µm across) with the cell’s segmentation painted on, drawn in the page. Can be switched off. | Quick ultrastructural check (nuclear shape, chromatin, cytoplasm) without opening the viewer. |
| B9 | **Cell history** | Chronological record of every identification, vote and proposal for the cell, with author and time. An earlier classification can be restored. | Audit trail for every label; curation is reversible. |
| B10 | **Organelles on the card** | Expandable list of the cell’s reported and traced structures, numbered per kind, each with a button that opens the viewer at it with its outline. | Shows what has been annotated in a cell and gives direct access to each item. |
| C1 | **Show in 3D** | In-page 3D view of the cell mesh (see-through) with the nucleus in blue, including community-proposed additional root IDs. | Checks morphology (processes, soma shape, relation to vessels) alongside ultrastructure. |
| C2 | **Nucleus volume** | Precomputed volume of the detected nucleus. | Nuclear size as a classification feature and for group comparisons. |
| C3 | **Cell volume** | Computes the volume of the cell’s mesh in the browser. The result is saved, and shown on the button (“Recalculate ≈ N µm³”) when already known. | Cell-size measurements across types; can be recomputed after segments are added. |
| C4 | **Mesh download (GLB) or + VR** | Downloads the cell mesh as a GLB file directly in the browser, with orientation corrected. +VR collects cells to show in a VR compatible environment. | Input for Blender, VR and 3D printing, and for figures. |
| C5 | **PowerPoint turntable** | Exports a slide with a rotating 3D model of the cell. | Teaching and presentations. |
| C6 | **Volume of traced structures** | Cavalieri estimate from traced contours, with a trapezoidal lower bound; inner contours are subtracted as holes. | Volumes of organelles and cells that have no automated segmentation. |
| C7 | **2D shape descriptors** | Per contour: circularity, aspect ratio, roundness and solidity, using ImageJ’s definitions. | Values are directly comparable with FIJI ROI measurements, e.g. lysosome shape in diseased versus healthy tissue. |
| C8 | **3D shape descriptors** | Per traced structure: sphericity (Wadell and MorphoLibJ conventions, both reported), elongation, flatness and solidity, in physical units. | Separates elongated from flattened organelles that look alike in a single section. |
| C9 | **Organelle distances** | Centroid and surface distances from each organelle to the nucleus and to its nearest sibling organelle. Saved with the measurement time and the number of siblings at that time. | Perinuclear distribution and clustering analyses, graphable from the stored table. |
| D1 | **Guided identification tree** | Step-by-step questions on ultrastructural and 3D features that lead to a cell type, with the criteria cited. | Standardises criteria across annotators and trains new users. |
| D2 | **Identification reports** | Each identification is recorded as a confirmation of, discrepancy with, or new identification relative to the model prediction, with certainty and comments. | Quantifies agreement with the model per cell type. |
| D3 | **Voting and consensus** | Users vote on identity claims and on root-ID proposals; the winning community name is shown on the card. | Consensus identities from independent annotators. |
| D4 | **New cell / Not a nucleus** | Reports cells missed by the nucleus detection, and flags detections that are not nuclei. | Corrects false negatives and false positives of the automated detection. |
| D5 | **Merged-nucleus split** | Reports the individual cells within one detection that contains several nuclei; each sub-cell gets its own entry, root ID and reports. | Recovers cells hidden inside merged detections. |
| D6 | **Root-ID proposals** | Proposes additional segments that belong to the cell. They are included in the 3D view, volume and viewer links. | Community proofreading of split segmentations. |
| D7 | **Master cell list** | One row per cell in a Google Sheet, updated with each report: current identity, layer, organelles, measurements and root IDs. | A single, analysis-ready table of the curated dataset. |
| E1 | **Organelle reporting** | Point or line annotations of 61 structure kinds in 13 groups (e.g. nucleus, centriole and cilium, endomembrane system, vesicles, cytoskeleton, junctions, vascular and perivascular contacts, synaptic structures, extracellular matrix, pathology). Can be logged from any identity card. | Systematic, cell-linked annotation of ultrastructure. Lines capture the extent of elongated structures such as primary cilia and nucleoplasmic reticulum. |
| E2 | **Bulk organelle annotation** | Paste a Neuroglancer link with markers of one kind. Each marker is assigned to its cell by reading the segmentation (root ID and nucleus ID) at that voxel, in the browser. | Fast annotation of one structure across many cells. |
| E3 | **“What the… is this?”** | A structure kind for flagging unknown structures for expert review. | Collects ambiguous cases for adjudication; flagged points can be filtered and viewed together. |
| F1 | **Tracing pad** | Draws an EM section in the page, optionally with the cell’s segmentation. Contours are placed point by point or drawn freehand with a pen or stylus; pan, zoom and section controls; rotating usage tips. AI tools for suggesting, outlining and following structures are available on the same pad (J1–J6). | Manual segmentation of structures the automated segmentation does not cover. |
| F2 | **Import from Neuroglancer** | Reads polygon, polyline and line annotations from a pasted link. Lines that nearly meet become one contour, and each annotation layer becomes its own structure. A 3D preview is shown before saving. | Reuses tracings made in Neuroglancer. |
| F3 | **Structure types and numbering** | The kind is chosen from the organelle list, plus whole cell and nucleus. Several structures of one kind are numbered 1…n and coloured apart. | Same vocabulary for annotation and segmentation; individual organelles stay identifiable. |
| F4 | **Contour editing** | Contours inside contours become holes. Contours can be interpolated in z (Catmull–Rom), and redundant points and sections removed within a set tolerance in nanometres. | Accurate geometry with fewer points; smoother 3D reconstructions. |
| F5 | **Live 3D preview** | The lofted surface updates while drawing, with the cell and nucleus shown see-through. | Immediate visual check of the reconstruction. |
| F6 | **Shared, versioned tracings** | Tracings belong to the dataset and can be extended by others. Every change is saved as a version with its author; unchanged structures are not re-saved. Drafts follow the account. Outlining a whole cell and naming its type counts as an identification. | Collaborative segmentation with attribution and full history. |
| F7 | **Open tracings in the viewer** | Opens tracings in Neuroglancer as polylines, centred on the cell. When a link is too long for the browser it opens in Spelunker or offers the JSON state instead. | Checks tracings against the full dataset. |
| G1 | **Identity filters** | Selects cells by type (every type listed, with counts, dimmed at zero), including cells with a traced whole cell or nucleus. | Defines the population to study. |
| G2 | **Spatial filters** | Vessel proximity and class, distance to glia limitans and white-matter border, V1 column membership, cortical layer, and one or more region-of-interest boxes. | Anatomically targeted populations, e.g. layer 1 astrocytes within 5 µm of a capillary. |
| G3 | **Organelle filters** | Presence or absence of reported or traced structures of selected kinds. | e.g. glia-limitans astrocytes with nucleoplasmic reticulum type II. |
| G4 | **Preview and count** | Lists the matches and states how many there are. | Immediate sample size before exporting. |
| G5 | **Excel export** | Exports matched cells with IDs, coordinates, identity, layer, distances, region box, cell history and measurements. | Direct input for statistics in Excel, R or Python. |
| G6 | **Open all matches in Neuroglancer** | One viewer state with all matched cells. Optional layers: EM, vasculature, V1 column boundary, capillary skeletons coloured by branch order (Spelunker), organelle points, centriole and cilium vectors, traced outlines. Offers the JSON if the state is too large. | 3D overview of a population in its tissue context. |
| G7 | **Step through matches** | Visits the matches one by one in a fixed, resumable order with a checked/remaining counter; each loads the identity card. | Exhaustive review of a defined population without missing cells. |
| G8 | **Blender export** | Generates a Google Colab notebook for a chosen box that builds a Blender scene: EM sections, segmentation overlay, cell meshes, nuclei, vasculature and traced structures, animated, with scale bar, box centre and cell identities. | Publication figures and videos from the same selection. |
| H1 | **Connectivity panel** | For a neuron, reads its synapses from the MICrONS synapse table (CAVE token required) and plots input/output synapse depth, synapse versus target soma depth, and target cell-type distribution. | Relates a cell’s identity to its wiring. |
| H2 | **Statistics dashboard** | Charts of cell types, layers, organelles and community activity, refreshed from the live data. | Monitors dataset composition and annotation progress. |
| I1 | **Google sign-in** | Credits every report to an account; warns before sending when signed out and detects an expired sign-in. | Every data point is attributable. |
| I2 | **Points and levels** | Points per action with per-cell caps and a bonus when later users agree with the first claimant; levels from Novice (0) to Godlike (100,000 points). | Motivation, and a basis for weighting annotators by track record. |
| I3 | **Combined leaderboard** | Ranks contributors across all tools in the family. | Visible recognition of contributors. |
| I4 | **Download your work** | Exports the user’s own reports. | Personal record and data reuse. |
| I5 | **References tab** | The literature behind the identification criteria. | Makes the criteria traceable. |
| I6 | **Build stamp** | Shows the build date and time of the page. | Records which version produced a result. |
| J1 | **Organelle suggestions** | A neural network running in the browser (ONNX, WebGPU or WASM; no server) marks organelles and structures on the EM section, e.g. lysosomes, mitochondria, Golgi, myelin, synapses, nuclear envelope and basement membrane. It reads the section and the sections 80 nm above and below. Classes that fail a precision test are hidden. | Speeds up annotation and points users to structures they might miss. Held-out validation: mean F1 0.80; lysosome precision/recall 0.98/0.98, mitochondria 0.92/0.95, Golgi 0.95/0.92. |
| J2 | **Correcting the machine learning** | Clicking a mark or a missed structure records it as right, not an organelle, an imaging artifact, or a different kind. Corrections are saved as attributed reports, separate from cell annotations. | Every correction becomes a training label, so the model improves with use; knife marks and folds are learned as their own class. |
| J3 | **AI outline** | One click inside an organelle returns its outline on that section (U-Net, 2 µm window at 8 nm). Outlines can be extended, cut or deleted with the pen. | Turns tracing an organelle from many clicks into one, while the user stays in control of the final contour. |
| J4 | **Nucleus outline model** | A second model (12 µm window at 32 nm) outlines nuclei. It was trained on the MICrONS nucleus segmentation plus hand-traced nuclei, and resamples the EM so it can run on other datasets. | Nuclear segmentation in datasets that have none, such as Lee16 and vCLEM. |
| J5 | **Follow through sections** | Carries an outline up or down the stack section by section. It stops when the area or position changes abruptly, and skips damaged sections (knife marks, missing tiles). | 3D reconstruction of an organelle or nucleus from one starting click. |
| J6 | **Machine learning (ML) -weighted gap filling** | Fills the sections between hand-drawn outlines 2–25 sections apart by blending interpolation with the ML prediction. The ML’s weight is set by how well it matched the user’s own outlines on the neighbouring sections, and the user’s outlines are never changed. | Sparse hand tracing (e.g. every 3–5 sections) gives a full reconstruction. The ML contributes only where it has proven reliable on that structure. |
| J7 | **Retrainable models** | Google Colab notebooks retrain the models from the community’s organelle points, tracings and corrections, combined with public ground truth: pinky100 mitochondria, MICrONS nuclei, MitoEM-H and CellMap. Each dataset can have its own models and otherwise uses µJump’s. | The models grow with the annotations and can be adapted to new datasets without changes to the tool. |

### **Supplementary Table S2. Datasets in ωJump**

*Sixty-two publicly released volumes from four repositories. ωJump streams each volume from the repository that publishes it and redistributes no image data. Volume identifiers are those used by the host repository and by the tool. Reference numbers index the list below the table.*

| **#** | **Volume identifier** | **Tissue** | **Species** | **Repository** | **Ref.** |
| --- | --- | --- | --- | --- | --- |
| 1 | jrc_hum-airway-14953vc | airway epithelium | *human* | OpenOrganelle | 1 |
| 2 | jrc_mus-choroid-plexus-3 | choroid plexus | *mouse* | OpenOrganelle | 1 |
| 3 | jrc_mus-duodenum-1a | duodenum | *mouse* | OpenOrganelle | 1 |
| 4 | jrc_mus-guard-hair-follicle | guard hair follicle | *mouse* | OpenOrganelle | 1 |
| 5 | jrc_mus-heart-1 | heart | *mouse* | OpenOrganelle | 1 |
| 6 | jrc_mus-kidney | kidney | *mouse* | OpenOrganelle | 1 |
| 7 | jrc_mus-kidney-2 | kidney (kidney-2) | *mouse* | OpenOrganelle | 1 |
| 8 | jrc_mus-liver | liver | *mouse* | OpenOrganelle | 1 |
| 9 | jrc_mus-liver-zon-1 | liver (zonation, zon-1) | *mouse* | OpenOrganelle | 1 |
| 10 | jrc_mus-liver-zon-2 | liver (zonation, zon-2) | *mouse* | OpenOrganelle | 1 |
| 11 | jrc_mus-lung-2a | lung | *mouse* | OpenOrganelle | 1 |
| 12 | jrc_mus-meissner-corpuscle-1 | Meissner corpuscle | *mouse* | OpenOrganelle | 1 |
| 13 | jrc_mus-pacinian-corpuscle | Pacinian corpuscle | *mouse* | OpenOrganelle | 1 |
| 14 | jrc_mus-pancreas-2 | pancreas (pancreas-2) | *mouse* | OpenOrganelle | 1 |
| 15 | jrc_mus-pancreas-4 | pancreas | *mouse* | OpenOrganelle | 1 |
| 16 | jrc_mus-skel-muscle-1 | skeletal muscle | *mouse* | OpenOrganelle | 1 |
| 17 | jrc_mus-skin-1 | skin | *mouse* | OpenOrganelle | 1 |
| 18 | jrc_mus-thymus-1 | thymus | *mouse* | OpenOrganelle | 1 |
| 19 | jrc_ut21-1413-003 | kidney (renal carcinoma) | *human* | OpenOrganelle | 1 |
| 20 | jrc_zf-cardiac-1 | heart | *zebrafish* | OpenOrganelle | 1 |
| 21 | wk-kasthuri2011 | cortex | *mouse* | WEBKNOSSOS | 2 |
| 22 | wk-cochlea | cochlea | *mouse* | WEBKNOSSOS | 3 |
| 23 | wk-morgan2020 | thalamus (dLGN) | *mouse* | WEBKNOSSOS | 4 |
| 24 | wk-bock11 | cortex (V1) + leptomeninges | *mouse* | WEBKNOSSOS | 5 |
| 25 | wk-k0563 | retina (k0563) | *mouse* | WEBKNOSSOS | 6 |
| 26 | wk-e2006 | retina (e2006) | *mouse* | WEBKNOSSOS | 7 |
| 27 | wk-k0725 | retina (k0725) | *mouse* | WEBKNOSSOS | 8 |
| 28 | wk-wanner16 | olfactory bulb | *zebrafish* | WEBKNOSSOS | 9 |
| 29 | wk-meissner-sham | Meissner corpuscle (sham, SBEM) | *mouse* | WEBKNOSSOS | 10 |
| 30 | wk-bloss18 | hippocampus | *mouse* | WEBKNOSSOS | 11 |
| 31 | wk-zf-vertebra | spinal cord | *zebrafish* | WEBKNOSSOS | 12 |
| 32 | wk-ex145-l4-s1 | cortex L4 (S1) | *mouse* | WEBKNOSSOS | 13 |
| 33 | wk-gour21-p7-l4 | cortex L4 (barrel, P7) | *mouse* | WEBKNOSSOS | 14 |
| 34 | wk-ex144-l23-s1 | cortex L2/3 (S1) | *mouse* | WEBKNOSSOS | 15 |
| 35 | wk-h5-l16-msem | cortex L1-6 (STG), MultiSEM | *human* | WEBKNOSSOS | 15 |
| 36 | wk-h5-l23-msem | cortex L2/3 (STG), MultiSEM | *human* | WEBKNOSSOS | 15 |
| 37 | wk-h5-l23-stg | cortex L2/3 (STG) | *human* | WEBKNOSSOS | 15 |
| 38 | wk-h6-l23-ifg | cortex L2/3 (IFG) | *human* | WEBKNOSSOS | 15 |
| 39 | wk-mk1-f6-l23 | cortex L2/3 (S1) | *macaque* | WEBKNOSSOS | 15 |
| 40 | wk-mk1-t2-stg | cortex L2/3 (STG) | *macaque* | WEBKNOSSOS | 15 |
| 41 | wk-denk2004 | cortex | *rat* | WEBKNOSSOS | 16 |
| 42 | wk-j0256 | HVC (song nucleus) | *zebra finch* | WEBKNOSSOS | 17 |
| 43 | wk-acc | cortex L2 (ACC) | *mouse* | WEBKNOSSOS | 18 |
| 44 | wk-karimi-lpta | cortex (LPtA) | *mouse* | WEBKNOSSOS | 18 |
| 45 | wk-karimi-s1 | cortex (S1) | *mouse* | WEBKNOSSOS | 18 |
| 46 | wk-ppc | cortex (PPC) | *mouse* | WEBKNOSSOS | 18 |
| 47 | wk-ppc2 | cortex L1-5 (PPC-2) | *mouse* | WEBKNOSSOS | 18 |
| 48 | wk-v2 | cortex L2/3 (V2) | *mouse* | WEBKNOSSOS | 18 |
| 49 | bd-ish21-retina-opl-em1 | retina, outer plexiform layer (EM1) | *mouse* | BossDB | 19 |
| 50 | bd-ish21-retina-opl-em2 | retina, outer plexiform layer (EM2) | *mouse* | BossDB | 19 |
| 51 | bd-wb21-nac-dopamine-cocaine | nucleus accumbens (dopamine axons, cocaine) | *mouse* | BossDB | 20 |
| 52 | bd-wb23-mouse-v1-l23-p105 | cortex L2/3 (V1, P105) | *mouse* | BossDB | 21 |
| 53 | bd-wb23-mouse-v1-l23-p523 | cortex L2/3 (V1, P523) | *mouse* | BossDB | 21 |
| 54 | bd-wb23-mouse-v1-l4-p105 | cortex L4 (V1, P105) | *mouse* | BossDB | 21 |
| 55 | bd-wb23-mouse-v1-l4-p14 | cortex L4 (V1, P14) | *mouse* | BossDB | 21 |
| 56 | bd-wb23-mouse-v1-l4-p523 | cortex L4 (V1, P523) | *mouse* | BossDB | 21 |
| 57 | bd-wb23-mouse-v1-l4-p6 | cortex L4 (V1, P6) | *mouse* | BossDB | 21 |
| 58 | bd-wb23-primate-v1-l23-p3000 | cortex L2/3 (V1, P3000) | *macaque* | BossDB | 21 |
| 59 | bd-wb23-primate-v1-l23-p7 | cortex L2/3 (V1, P7) | *macaque* | BossDB | 21 |
| 60 | bd-wb23-primate-v1-l23-p75 | cortex L2/3 (V1, P75) | *macaque* | BossDB | 21 |
| 61 | bd-wb23-primate-v1-l4-p3000 | cortex L4 (V1, P3000) | *macaque* | BossDB | 21 |
| 62 | ew2-stroeh-retina | retina (Eyewire II) | *mouse* | Eyewire II | 22 |
